# Genetics of mouse behavioral and peripheral neural responses to sucrose

**DOI:** 10.1101/2020.08.19.253989

**Authors:** Cailu Lin, Masashi Inoue, Xia Li, Natalia P. Bosak, Yutaka Ishiwatari, Michael G. Tordoff, Gary K. Beauchamp, Alexander A. Bachmanov, Danielle R. Reed

## Abstract

Mice of the C57BL/6ByJ (B6) strain have higher consumption of, and stronger peripheral neural responses to, sucrose solution than do mice of the 129P3/J (129) strain. To identify quantitative trait loci (QTLs) responsible for this strain difference and evaluate the contribution of peripheral taste responsiveness to individual differences in sucrose intake, we produced an intercross (F_2_) of 627 mice, measured their sucrose consumption in two-bottle choice tests, recorded the electrophysiological activity of the chorda tympani nerve elicited by sucrose in a subset of F_2_ mice, and genotyped the mice with DNA markers distributed in every mouse chromosome. We confirmed a sucrose consumption QTL (*Scon2, or Sac*) on mouse chromosome (Chr) 4, harboring the *Tas1r3* gene, which encodes the sweet taste receptor subunit T1R3 and affects both behavioral and neural responses to sucrose. For sucrose consumption, we also detected five new main-effect QTLs *Scon6* (Chr2), *Scon7* (Chr5), *Scon8* (Chr8), *Scon3* (Chr9) and a sex-specific QTL *Scon9* (Chr15), and an interacting QTL pair *Scon4* (Chr1) *and Scon3* (Chr9). No additional QTLs for the taste nerve responses to sucrose were detected besides the previously known one on Chr4 (*Scon2)*. Identification of the causal genes and variants for these sucrose consumption QTLs may point to novel mechanisms beyond peripheral taste sensitivity that could be harnessed to control obesity and diabetes.

## Introduction

Inbred mouse strains C57BL/6ByJ (B6) and 129P3/J (129) differ in sucrose consumption (Bachmanov, Reed et al. 1996, Bachmanov, Tordoff et al. 1996, Bachmanov, Reed et al. 1997, Bachmanov, Tordoff et al. 2001, Inoue, Reed et al. 2004, Sclafani 2004, Sclafani and Ackroff 2004, Inoue, Glendinning et al. 2007, de Araujo, Oliveira-Maia et al. 2008, Bachmanov, Bosak et al. 2011), with heritability ranging from 36% to 78%, depending on the measure of consumption used. There is a high heritability for sucrose intake but relatively low heritability for sucrose preference score owing to a ceiling effect (i.e., preference scores are nearly 100% for concentrated sucrose). The heritability of sucrose intake is partially determined by a genetic locus (*Scon2*, or *Sac*) on distal Chr4, which corresponds to the “sweet taste” gene (*Tas1r3*) coding for the T1R3 protein (Bachmanov, Li et al. 2001, Li, Staszewski et al. 2002, Reed, Li et al. 2004). Allelic variation in this gene affects consumption of sucrose and the other sweeteners (Inoue, Reed et al. 2004, Reed, Li et al. 2004, Inoue, Glendinning et al. 2007).

However, the *Scon2*/*Sac* locus explains only a part of the phenotypic variation in sucrose consumption. Knocking out the *Tas1r3* gene eliminates most behavioral responses to some sweeteners (Damak, Rong et al. 2003, Zhao, Zhang et al. 2003), but these mice still prefer concentrated sucrose, which suggests that genetic loci other than *Scon2*/*Sac* affect sucrose intake (Damak, Rong et al. 2003, Zhao, Zhang et al. 2003).

It is unknown whether these additional genetic loci act via peripheral taste mechanisms or postoral mechanisms (for example, by influencing sucrose metabolism). To assess whether genetic loci affect the peripheral taste mechanisms, investigators measure responses in the chorda tympani gustatory nerve to oral stimulation with sucrose (Ninomiya, Kajiura et al. 1993, Bachmanov, Reed et al. 1997, Inoue, McCaughey et al. 2001, Damak, Rong et al. 2003, Danilova and Hellekant 2003).

To detect novel quantitative trait loci (QTLs) that affect sucrose consumption (measured as sucrose solution intakes and preference scores in two-bottle choice tests), we intercrossed the B6 and 129 strains and conducted linkage analyses of the F_2_ mice; neural responses to sucrose were recorded in a subset of these F2 mice. To describe the genetic architecture of behavioral and neural responses to sucrose, we included epistasis and sex-specific effects in our statistical analysis strategy. We detected seven QTLs and named them sucrose consumption QTL 2-4, 6-9 (symbols *Scon2-4, 6-9*), which are mapped to Chr4, 9, 1, 2, 5, 8 and 15, respectively (*Scon1* and *Scon5* QTLs were detected in previous studies). We considered linkages for intakes and preference scores across sucrose concentrations as the same QTL if confidence intervals for different phenotypic measures overlapped and QTLs with the same effect direction. Our previous studies (Bachmanov, Li et al. 2001, Inoue, Reed et al. 2004, Reed, Li et al. 2004, Inoue, Glendinning et al. 2007) have shown that *Scon2, Sac* (saccharin preference locus) and *Tas1r3* (taste receptor, type 1, member 3 gene) are identical, and so here we use *Scon2* to describe linkages to the distal Chr4 (154.4-156.5 Mb) for 120 mM and 300 mM sucrose intakes and preferences as well as neural responses to sucrose (100-1000 mM).

## Materials and methods

### Animals and breeding

The B6 (stock no. 001139) and 129 (stock no. 000690) inbred mice were obtained from The Jackson Laboratory (Bar Harbor, ME, USA) and intercrossed to produce F_1_ and F_2_ hybrids in the Animal Facility of Monell Chemical Senses Center, located in Philadelphia, Pennsylvania (USA). Pups were weaned at 21-30 days of age and reared in groups of the same sex. The mice were housed in a temperature-controlled vivarium at 23°C on a 12/12-hr light-dark cycle, with lights off at 7 pm, barring unusual circumstances (e.g., power outages), and had ad lib access to tap water and food (Rodent Diet 8604, Harlan Teklad, Madison, WI, USA). We bred F_2_ mice for two separate experiments (**S1 Table**). All animal procedures in this study were approved by the Institutional Care and Use Committee of the Monell Chemical Senses Center.

### Phenotyping

#### Two-bottle taste preference tests

All F_2_ mice were tested for their intake and preference for sucrose and other solutions using two-bottle taste preference tests. Mice from the parental B6 and 129 strains (20 of each strain) were also included in the two-bottle tests in Experiment 2 (described below).

Procedure details for the two-bottle taste test have been described elsewhere (Tordoff MG 2001, Bachmanov, Reed et al. 2002, Bachmanov, Reed et al. 2002, Bachmanov, Reed et al. 2002). In brief, individually caged mice were presented with one tube containing a taste solution in deionized water and another tube containing deionized water only. Daily measurements were made in the middle of the light period by reading fluid volume to the nearest 0.1 ml. The positions of the tubes were switched every day to control for positional preferences (some mice prefer to drink from one side regardless of the contents of the tube). For 2 days before testing, mice were offered both tubes with deionized water.

Here, we present data for 96-hr (4-day) two-bottle tests with 3, 120, and 300 mM sucrose obtained in two experiments. In addition to sucrose, mice were also tested with other solutions (data are not shown). In **Experiment 1** (F_2_ males only) mice were offered the following taste solutions in this order: 300 and 75 mM NaCl, 120 mM sucrose, 0.1 mM citric acid, 10% ethanol, and 0.03 mM quinine hydrochloride. In **Experiment 2** (B6, 129, and F_2_; females and males) mice were offered the following taste solutions in this order: 30 mM glycine, 30 mM D-phenylalanine, 20 and 1 mM saccharin, 120, 300, 3 mM sucrose, 1 and 300 mM monosodium salt of L-glutamic acid (MSG), 3% and 10% ethanol, and 0.03 mM quinine hydrochloride. We purchased all taste compounds from Sigma Chemical Co. (St. Louis, MO, USA), except the ethanol, which was purchased from Pharmco Products (Brookfield, CT, USA).

Each concentration of a solution was presented for 4 days (Tordoff and Bachmanov 2002). Between tests of each taste compound, mice had water to drink from both tubes for at least 2 days to reduce potential carryover effects. This procedure was also used with sucrose between 300 and 3 mM (but not for other concentrations of sucrose and within other compounds). No measures of water intakes were available for Experiment 2, so we used total fluid intake of quinine hydrochloride (sum of water intake and quinine solution intake) as mouse water intakes, because almost all of mice avoid the quinine solution, and thus the measure of total fluid intake of quinine solution approximately equals water intake.

#### Electrophysiology

Electrophysiological experiments were conducted with a subset of 84 F_2_ mice selected from Experiments 1 and 2 and an additional F_2_ cross (Inoue, Reed et al. 2004) (**S1 Table**). The activity of the whole chorda tympani nerve in response to lingual application of taste solutions was recorded electrophysiologically for the following stimuli for each group of mice separately: for mice from Experiment 1 (19 males), 10, 30, 100, 300, and 1000 mM sucrose; for mice from Experiment 2 (58 mice: 29 males, 29 females), 3, 10, 30, 100, 300, 500, and 1000 mM sucrose; for the 7 additional F_2_ mice (2 males, 5 females), 500 mM sucrose. For all mice, 100 mM NH_4_Cl solution was presented at regular intervals to serve as a reference stimulus. During chemical stimulation of the tongue, the test solutions flowed for 30 sec (Inoue, Li et al. 2001, Inoue, McCaughey et al. 2001, Inoue, Reed et al. 2004). Between taste stimuli, the tongue was rinsed with deionized water for at least 1 min to offset carryover effects. The magnitude of the integrated response at 20 sec after stimulus onset was measured and expressed as a proportion of the average of the previous and following responses to 100 mM NH_4_Cl (Inoue, Li et al. 2001, Inoue, McCaughey et al. 2001).

#### Body weight and composition

Body weights were measured before and after each taste compound was offered in the two-bottle taste preference tests. Body weight did not change appreciatively during the 4-day taste tests, so we averaged the before and after body weights for some analyses. In addition, we performed body composition analyses for all Experiment 2 mice by necropsy, weighing the epididymal and retroperitoneal adipose depots to the nearest 0.01 g and summing these values as a proxy for total amount of dissectible fat per mouse. At necropsy we also used a ruler (±1 mm precision) to measure body length from the base of the teeth to the anus.

### Genotyping

Genomic DNA was extracted and purified from mouse tails by a sodium hydroxide method or by proteinase K digestion followed by high-salt precipitation (Gentra/Qiagen, Valencia, CA, USA). In total, 627 mice were genotyped with 429 polymorphic markers (see **S2 Table**), with control DNA samples from the F_1_ 129 and B6 inbred strains. Genotyping was conducted in two steps to achieve an average distance of 5.9 Mb between markers and no gap greater than 40 Mb. First, simple-sequence repeat markers known to be polymorphic between the parental strains were genotyped to cover all 19 autosomes and the X chromosome (simple sequence-length polymorphisms). Fluorescently labeled microsatellite primers were amplified by PCR, and the products were scanned by an ABI 3100 capillary sequencer (Applied Biosystems, Foster City, CA, USA). Second, single-nucleotide polymorphisms (SNPs) were added to fill gaps (KBiosciences, Herts, UK). A few SNPs were also genotyped in our laboratory, using primers and fluorescently labeled probes designed to discriminate alleles using an ABI Prism 7000 real-time PCR system (ABI Assay-by-Design, Applied Biosystems, Foster City, CA, USA). In a few cases, we typed an SNP marker and a simple sequence-length polymorphism marker that were adjacent, in nearly identical physical locations, and pooled these data into a single maker for linkage analysis. In addition, genotypes associated with coat and eye color were included in the linkage analysis.

### Data analyses

#### Taste solution intake and preference

For each concentration of taste solution, individual average solution and water intakes were calculated based on daily intake values. Preference scores were calculated for each mouse as the ratio of the average daily solution intake to average daily total fluid (solution + water) intake, in percent. Correlations among taste intakes and preferences for all solutions were evaluated using Pearson’s correlation coefficients. For correlational analyses among different taste solutions, we imputed missing data using an algorithm implemented in the MICE package (Multivariate Imputation by Chained Equations) (Resche-Rigon and White 2018) (**S1 Figure**).

#### Sex, genotype, and dominance effects

We pooled the data (129, B6, and F_2_) and tested whether males and females differed by sex and genotype using a two-way ANOVA with sex and genotype as fixed factors. We tested for dominance effects using with a *t*-test to compare the mean trait values in the F_2_ mice relative to the mean values from the parental strains (Bachmanov, Reed et al. 1996).

#### Preparing data for linkage analysis

Distributions (**S2 Figure**) and sex differences and normality (**S3 Table**) of the sucrose intakes and preferences of F_2_ mice and mice from each parental strain were analyzed using the R package *fitdistrplus* (Delignette-Muller).

To assess whether data need to be adjusted (standardized), we used observed (unadjusted) data to evaluate the effect of covariates (habitual water intake, sex, age, body fat, body length, and body weight) on sucrose intake and preference and found that habitual water intake, age, and sex were bona fide covariates but that body weight, body length, and body composition were not. For age, the range of age differed between Experiments 1 and 2, but the ages were similar within each experiment, and therefore standardizing sucrose intake and preference within each experiment eliminated the variation due to age. Thus, for the linkage analyses described below, we used data that were adjusted (standardized) separately for each sucrose concentration tested and for each index (intakes and preferences): for intakes, we calculated standardized residuals (residual values of the sucrose intake relative to habitual water intake, standardized within sex), and for preference scores, we calculated standardized scores (sucrose preference scores standardized within sex and experiment, if applicable).

We conducted similar analysis for the electrophysiology data. There were effects of sex but no other *bona fide* covariates. Therefore, we standardized the data within sex and experiment and used these adjusted (standardized) data for linkage analyses described below.

#### Linkage analyses

We first screened the genotype data for errors by determining whether genotypes were compatible with the preexisting haplotypes; genotypes such as those that created double recombinants were re-assayed.

Genome-wide scans of the F_2_ mice for the sucrose consumption and electrophysiological data were conducted using markers from 19 mouse autosomes and the X chromosome. The degree of linkage between individual traits and genotypes was computed using algorithms implemented by the R/QTL1.42 package of R (Broman, Wu et al. 2003). Genotype probabilities and genotype errors were estimated using the *calc*.*genoprob* function, and the missing genotypes were imputed using the *sim*.*geno* function. Interval mapping by maximum likelihood estimation (EM algorithm) was conducted to screen for main effect QTLs using the *scanone* function. The significance of each marker regression result was established by comparison with 1000 permutations of the observed data using the *n*.*perm* function. The main-effect confidence intervals were defined by a drop of 1 LOD (logarithm of the odds) from peak linked marker and were calculated by applying the *lodint* function and expanded to the outside markers.

We also assessed sex-specific effects using R/QTL to calculate LOD scores for two models, “sex additive” and “sex additive with interactions” (Solberg, Baum et al. 2004), and we calculated the difference in LOD scores between the genome scan results obtained by including these two models separately using the *arithscan* function. To quantify sex-by-genotype interactions, we compared the fit of the two models using the difference in LOD scores as a metric (ΔLOD) and using a cutoff ΔLOD 2.0 as a criterion (Lin, Theodorides et al., Lin, Theodorides et al. 2013).

Two-dimensional genome scans were used to estimate marker pair interactions (epistasis) using the *scantwo* function. The LOD score cutoff for significant epistasis was determined by a comparison with 1000 permutation tests at the p<0.05 level and confirmed in a general linear model using marker pairs as a fixed factor and habitual water intake as a covariate.

To assess dominant/additive interactions between the QTL alleles, the trait mean for the 129/B6 heterozygotes for the marker at peak linkage was compared with the collapsed trait value for the 129/129 and B6/B6 homozygotes; these tests were conducted using planned comparisons.

#### Statistical models and programs

For all statistical models, we used a type 1 (sequential) sum-of-squares model, and when testing for group differences, we used Tukey’s HSD tests. We computed statistical tests with R (version 3.3.3) and RStudio (version 1.0.136) and graphed the results using either R or Prism 6 (version 6.05; GraphPad Software, La Jolla, CA, USA).

### Candidate gene analyses

We defined the QTL critical regions by 1-LOD drop from the peak linked marker and expanded them to the most adjacent outside marker. We merged regions for sucrose QTLs if they were on the same chromosome with overlapping confidence intervals. The resulting chromosome regions were used to evaluate candidate genes, as follows: *Scon6* @ chr2:100122595-128989913, *Scon2* @ chr4:154415509-1558633334, *Scon*7@ chr5:22747915-66662082, *Scon8* @ chr8:72545028-129008800, *Scon3* @ chr9:87885544-110762590, *Scon9* @ chr15: 40369844-71152967. For the critical region of *Scon4* on Chr1 (epistatic QTL), we defined the region using the peak linked marker *rs3714728* (9.0 Mb) with a 20-Mb flanking region (chr1:1-19023602). Within these regions, we listed the known genetic variants and evaluated them in several ways. First, we used information in the Mouse Genomes Project-Query SNPs, indels, or structural variations (Anonymous Mouse Genomes Project -Query SNPs, indels or SVs. 2011: Wellcome Trust Sanger Institute) to make the list. This database contains the results of a large-scale genome sequencing project of many inbred mouse strains, including the 129P2/OlaHsd and C57BL/6J strains (Keane, Goodstadt et al. 2011, Yalcin, Wong et al. 2011), which are closely related to the 129 and B6 parental inbred strains used in our study (Yang, Wang et al. 2011). From this list, we identified sequence variants with the potential to cause functional changes using the PROVEAN web server (Choi and Chan 2015). Next, using the Mouse Genome Data Viewer (Anonymous), we identified all genes residing within the chromosome critical regions and classified these candidate genes using the PANTHER system (Mi, Muruganujan et al. 2013).

Using results from the two-dimensional scan, we determined if there were candidate genes from the interacting QTLs *Scon3* and *Scon4* that had the same or similar biological function, reasoning that if the two genes have epistatic interaction, they likely act in the same pathway to regulate sucrose intake. To that end, we identified interacting gene pairs and probed their protein-protein associations by searching the STRING database (von Mering, Jensen et al. 2005, York, Reineke et al.). We exported gene pairs for protein-protein interactions with association scores > 0.9, which were computed by benchmarking them against the training set (von Mering, Jensen et al. 2005), and identified those pairs that fit our criterion (confidence score = 0.9).

Finally, for the genes (*N*=2565) within the QTL confidence intervals, we used publicly available GEO data (Davis and Meltzer 2007) to compare patterns of these gene expression at three tissues (striatum, brown and white fat) between the 129S1/SvImJ and C57Bl/6J inbred strains, which are closely related to the 129 and B6 strains used in our study. For mouse striatum, we downloaded microarray data for three available samples from each strain (129P3/J and C57BL/6J; GSE7762 (Korostynski, Piechota et al. 2007)) and conducted gene expression analysis with the web tool GEO2R (http://www.ncbi.nlm.nih.gov/geo/geo2r). We calculated the distribution of fluorescence intensity values for the selected samples and then performed a linear model for microarray data (LIMMA) statistical test (Smyth 2005) after applying log transformation to the data. For fat, we downloaded bulk RNASeq data for four available samples from each strain (129S1/SvImJ and C57Bl/6J; GSE91067 (Raymond E Soccio 2017) and each tissue (brown and white fat) and transformed the transcript count data. We created a matrix for the two groups (129 vs B6) and fitted data with the genewise glmFit function of the edgeR package (Robinson, McCarthy et al. 2010); we conducted likelihood ratio tests for 129 vs B6 for brown and white fat tissue separately. The differentially expressed genes for the two types of gene profiling data sets were reported as one log_2_-fold change (corresponding to a twofold change) with an associated p-value corrected for false discovery (FDR<0.05) for multiple tests (Benjamini 1995).

## Results

### Phenotypes

#### Sucrose intake; preference in B6, 129, and F_2_ mice; and sex-specific hereditary model

On average, B6 mice drank significantly more sucrose and had significantly higher preferences for 3, 120, and 300 mM sucrose than did 129 mice (**Figure 1**; **S3 Figure**), with F_2_ mice falling between the means of the parental groups. Female mice drank more sucrose on average and had a higher preference for sucrose compared with male mice of the same genotype (**Figure 1**; **S3 Figure, S3 Table**). Significant interactions between sex and genotype were observed for 120 and 300 mM sucrose intake and preference but not for 3 mM sucrose (**S4 Table**). For 120 and 300 mM sucrose, all female mice regardless of genotype had similarly high sucrose intakes and preferences, but B6 males had higher sucrose intakes and preferences than did 129 males (**Figure 1**). The inheritance pattern was dominant (as measured by the significant difference between the observed mean of the F_2_ mice and the theoretical midparent mean) except for 3 and 300 mM sucrose preference in females (**S5 Table**).

**Figure 1.**
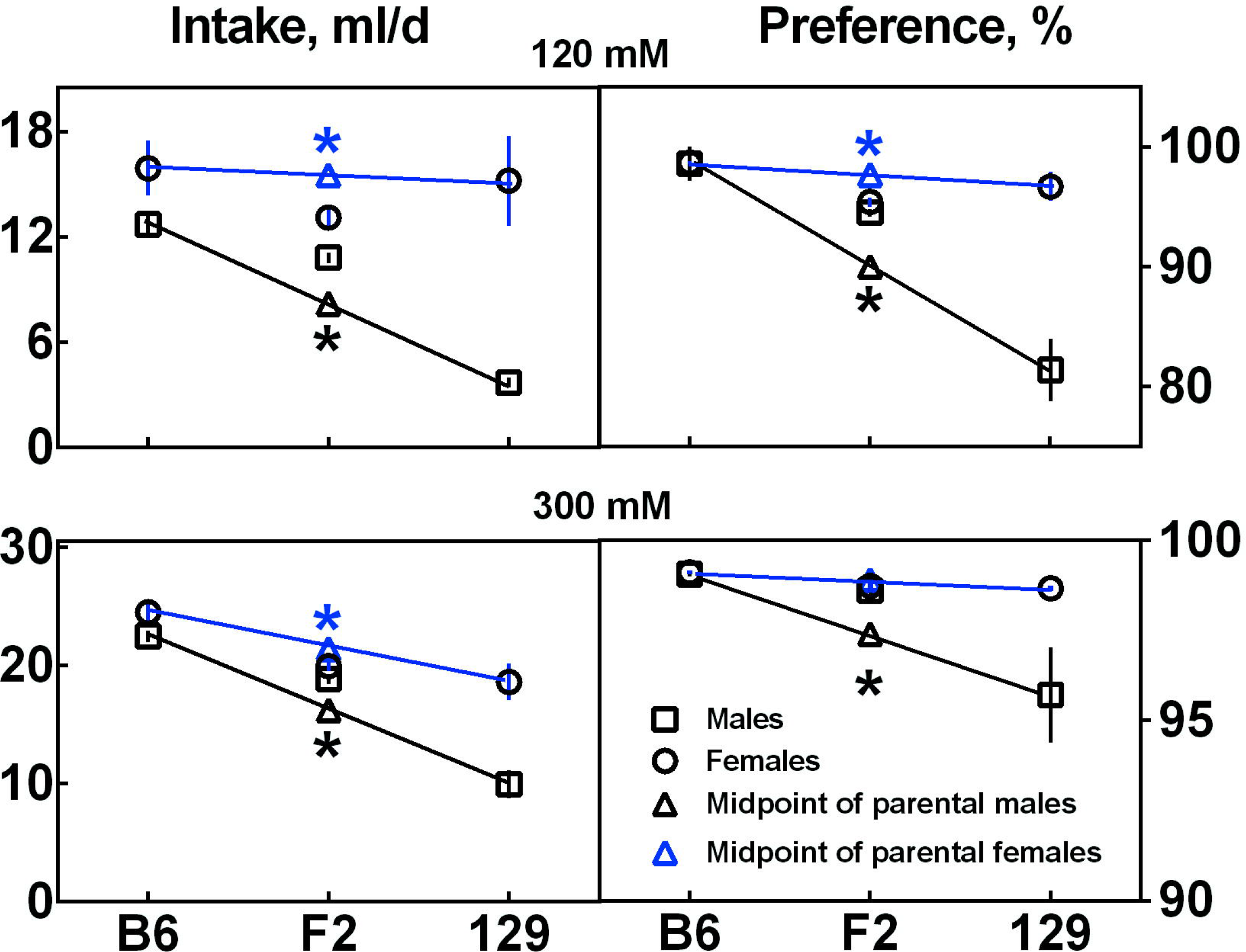
Dominance effects for sucrose intake and preference by sex. Data are mean and standard error for intake (left) and preference (right) of 120 mM (top) and 300 mM (bottom) sucrose solutions by male and female parental strains (B6, 129) and F_2_ mice. Values for the F_2_ mice (data points with center bar) are displayed at the parent midpoint to show the direction of dominance, if any. Asterisks indicate a dominant mode of inheritance (as defined by a significant difference between the F_2_ group average and the parental midpoint).

#### Correlations between indexes of consumption of taste solutions

In the F_2_ mice, sucrose intake correlated with sucrose preference score, but this relationship was weaker at the higher concentrations (**S6 Table;** observed data), most likely due to a ceiling effect, with truncated ranges of preference scores for concentrated sucrose (near 100%; **Figure 1**). Within sucrose intakes and preferences, traits also correlated among concentrations for intakes and preference scores (**S4 Figure**; imputed data).

#### Body size and composition

There were no significant correlations between sucrose intake and body weight, body length, or body fat in F_2_ mice (**S5A-F Figures**) except a weak correlation between 3 mM sucrose intake and body weight in females (*r*=0.15, *p*=0.02) that did not extend to the adjusted residual scores of sucrose intakes (*r*=0.09, *p*=0.19, **S5B Figure**). There were similar correlations between sucrose intake and body length in males (3 mM: *r*=0.15, *p*=0.02; 120 mM: *r*=0.16, *p*=0.02; 300 mM: *r*=0.16, *p*=0.02) that did not extend to the adjusted residual scores of sucrose intakes (**S5E Figure**). There was no correlation between sucrose preference and body weight, body fat, or body length (**S6A-F Figures**) except for a weak correlation between 3 mM sucrose preference and body length in females (*r*=0.17, *p*=0.01; **S6F Figure**).

#### Habitual water intake

There was a significant mouse strain effect on habitual water intake (*F*_(1, 654)_ = 18.9, *p*<0.0001, two-way ANOVA; **S4 Table**). In the F_2_ mice, sucrose intakes significantly correlated with habitual water intakes, but as expected, the adjusted residual scores of sucrose intakes did not correlate with habitual water intake(**S5G Figure**). There was no correlation between habitual water intakes and sucrose preferences, using either unadjusted or adjusted scores (**S6G Figure**).

#### Sex

In addition to the genotype effects, we observed significant sex and/or sex-by-genotype interaction effects on sucrose intake, especially for the concentrated sucrose solutions (120 and 300 mM; **S4 Table**). Significant sex effects on water intake and body weight were observed (**S4 Table**).

#### Age

When the two-bottle tests began in Experiment 1, the F_2_ mice were 66 ± 0.6 days old (range, 60-90 days); for Experiment 2, the F_2_ mice were 162.9 ± 1.2 days old (range, 117-204 days), and the B6 and 129 mice were 120 ± 3 days old (range, 90-147 days). At the time of conducting electrophysiological experiments, the F_2_ mice from Experiment 1 were 81 ± 0.6 days old (range, 75-105 days), and those from Experiment 2 were 279 ± 3 days old (range, 234-351 days). The 7 mice from the additional cross were adult (but the exact ages were not recorded). At the time of performing the body composition analyses, the F_2_ mice from Experiment 2 were 267 ± 1.5 days old. There was no significant correlation (*r*<0.1, *p*>0.05) between sucrose consumption (intake and preference) and age (**S5H** and **S6H Figures**), except a weak correlation between unadjusted but not adjusted 120 mM sucrose preference and age (unadjusted, *r*=0.09, *p*=0.02; adjusted, *r*=0.01, *p*=0.88; **S6H Figure**).

### Linkages

#### Main-effect QTLs

For 120 mM sucrose solution, chromosome mapping identified QTLs on Chr4 (*Scon2*) and Chr9 (*Scon3*) for both intake (*Scon2*: LOD = 27.1; *Scon3*: LOD = 5.3) and preference (*Scon2*: LOD = 59.2; *Scon3*: LOD = 4.1), with the B6 allele increasing the traits (**Figure 2**; **S7 Table**); because these QTLs for sucrose intake and preference were detected in the same chromosomal location with overlapping confidence intervals, we named them as the same QTL symbols (**S7 Table)**. There was a QTL on Chr5 (*Scon7*) for 120 sucrose preference (LOD = 4.1), with the 129 allele increasing the phenotype (**Figure 2**; **S7 Table**). The LOD scores of the peak marker, confidence intervals of these QTLs, and fractions of genetic variance they explain (4-90%) are summarized in the **S7 Table**.

**Figure 2.**
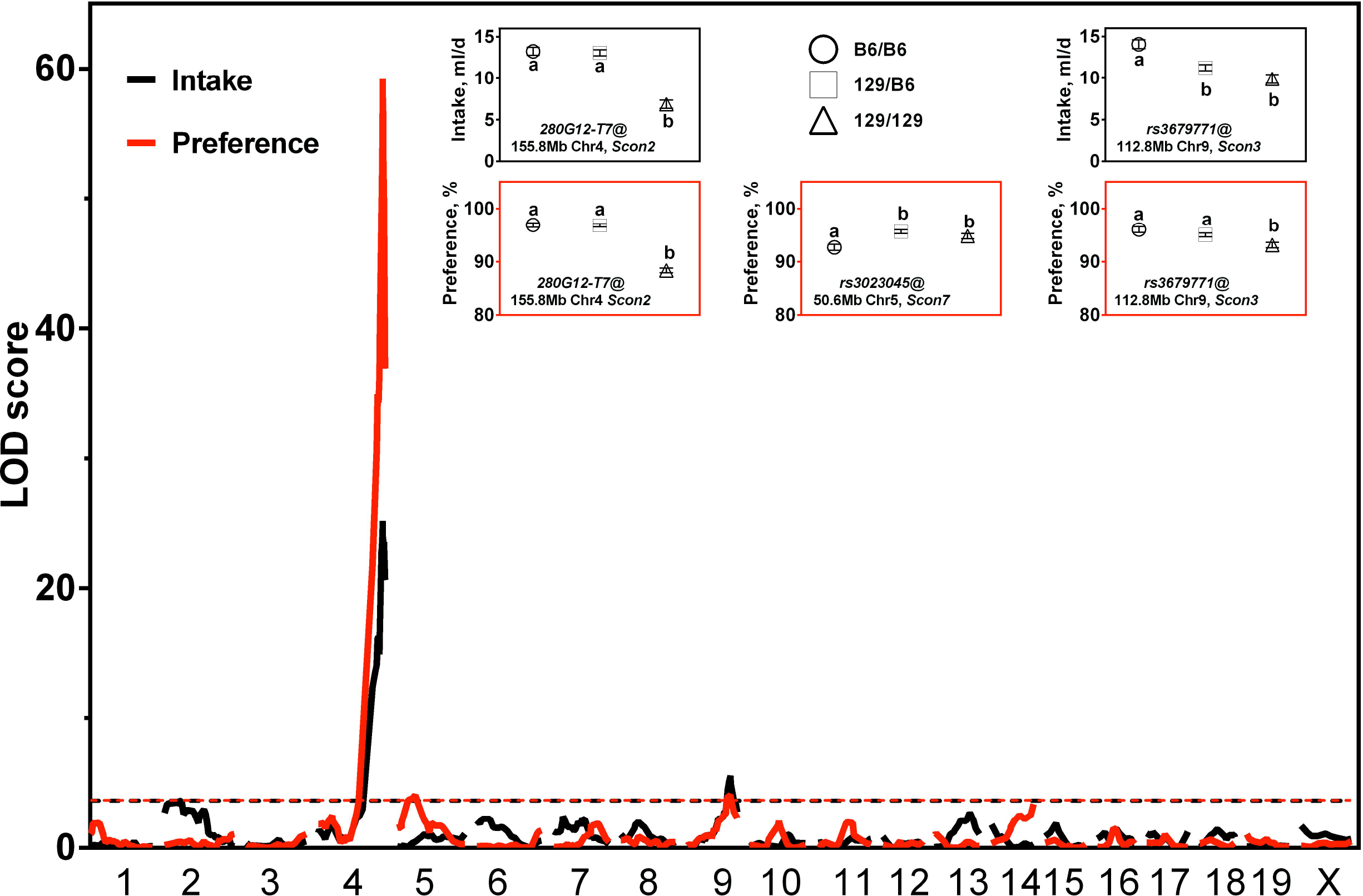
Genome-wide scan of main-effect QTLs for 120 mM sucrose intake and preference. The x-axis shows chromosome numbers and chromosomal locations (in Mb). The y-axis shows the logarithm of the odds (LOD) scores calculated for adjusted intakes and preferences. The horizontal dashed lines represent significant linkage thresholds that correspond to an LOD of 3.84 for intake (black) and 3.68 for preference (red). Boxed insets show least-square means (LSM) and standard errors (SE) for sucrose intakes (black frames) and preferences (red frames) of the F_2_ mice, grouped by genotype of the markers nearest to the linkage peak. Letters (a, b), when different, indicate statistically significant differences between genotypes (the results of post hoc tests in a general linear model with habitual water intake as a covariate).

For 300 mM sucrose solution, we identified four main effect QTLs for intakes (*Scon6, Scon2, Scon8 and Scon3* on Chr 2, 4, 8, and 9) and *Scon3* (on Chr9) for preference (**Figure 3**; **S7 Table**), with the B6 allele increasing the trait for all QTLs except for *Scon6*, for which the 129 allele increased the trait. The LOD scores of the peak marker, the confidence intervals of these QTLs, and fractions of genetic variance they explain (4-26%) are summarized in the **S7 Table**.

**Figure 3.**
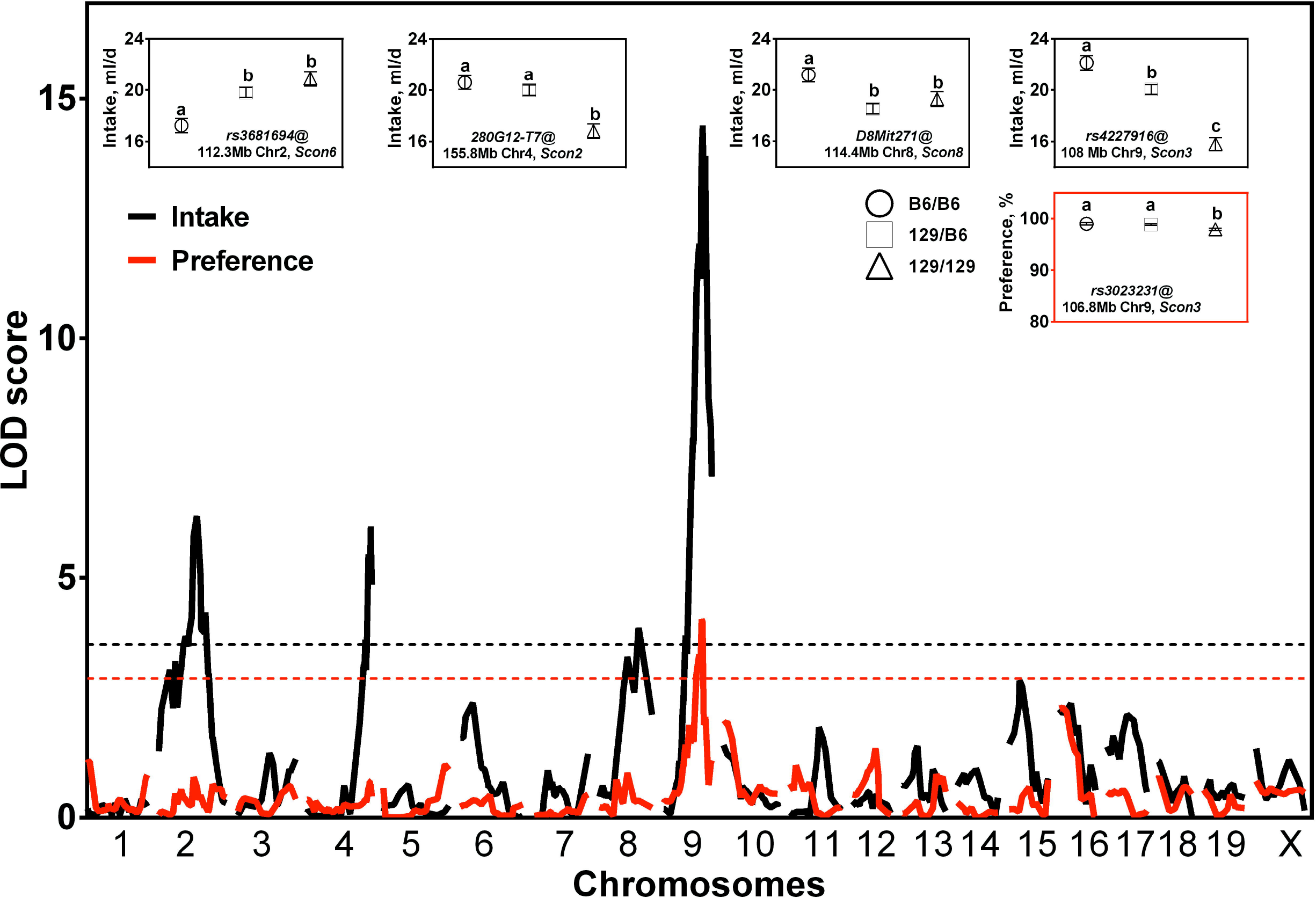
Genome-wide scan of main-effect QTLs for 300 mM sucrose intake and preference. The horizontal dashed lines represent significant linkage thresholds that correspond to an LOD of 3.61 for intake (black) and 2.90 for preference (red). Other details are the same as for **Figure 2**.

#### Sex-specific QTL

For 300 mM sucrose intake, there was a male-specific QTL (*Scon9)* on Chr15 with the peak LOD score of 4.73 at peak marker *rs4230721* (54.7 Mb), with a confidence interval of 40.3-71.1 Mb (between *rs6396894* and *rs3088560*); it explained 7% of the genetic variance in sucrose intake, with the B6 allele increasing the phenotype (**Figure 4**). This male-specific effect was confirmed in a general linear model using habitual water intake as a covariate (sex: *F*_(1,444)_=3.59, *p*=0.058; genotype of *rs4230721*: *F*_(2, 444)_=5.08, *p*=0.006, sex × genotype of *rs4230721*: *F*_(2,444)_ =4.05, *p*=0.018).

**Figure 4.**
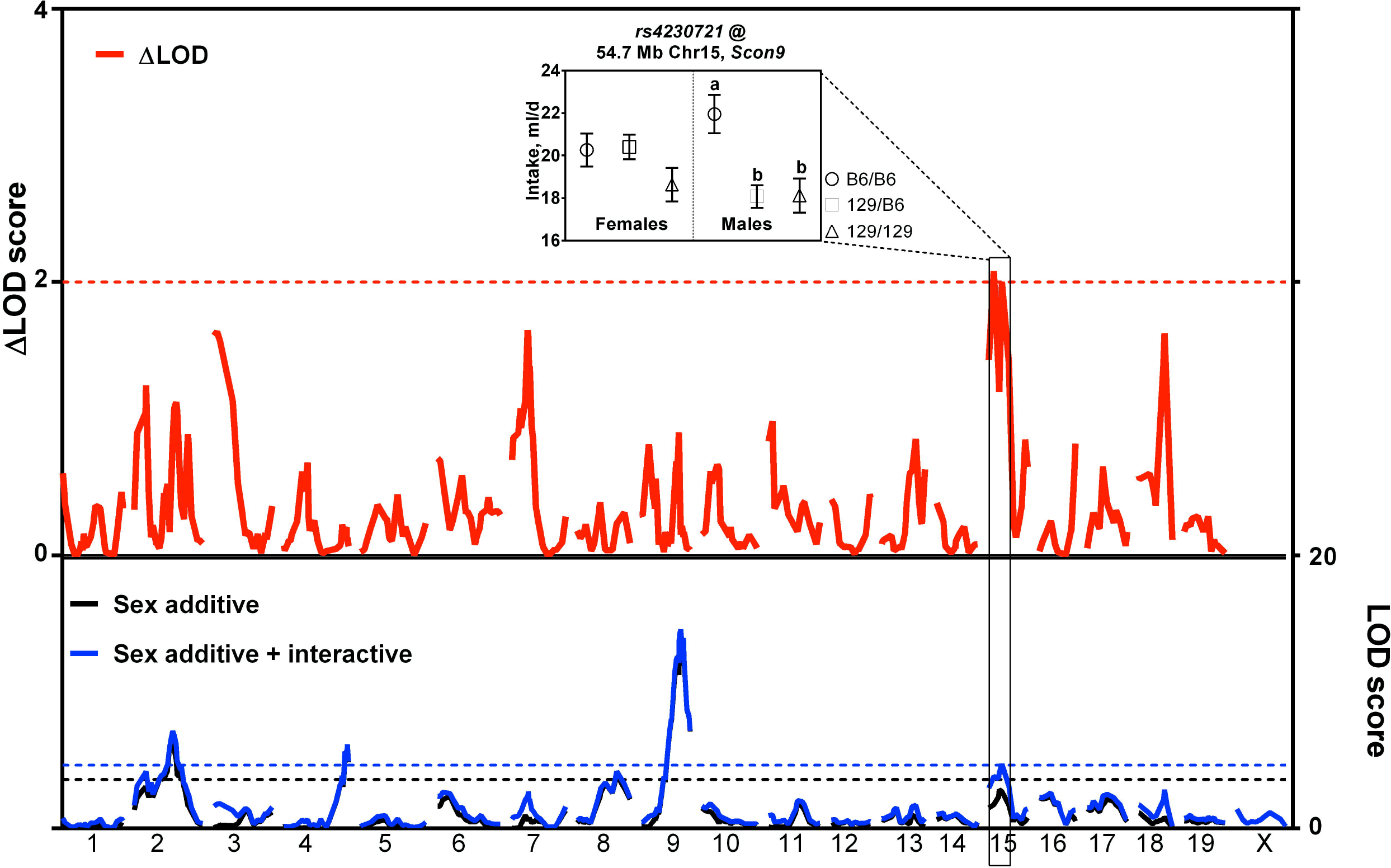
Genome-wide scan of sex-specific QTLs for 300 mM sucrose intake. **Bottom panel,** Plots of LOD scores (right y-axis) for genome scans of 300 mM sucrose with sex as additive covariate (black curve; black horizontal dashed line shows significant linkage thresholds that correspond to an LOD of 3.61) and sex as additive plus interactive covariate (blue curve; blue horizontal dashed line shows significant linkage thresholds that correspond to an LOD of 4.65). Significant linkages were detected on Chr2, 4, 8, and 9 using sex as additive covariate and on Chr2, 4, 8, 9, and 15 using sex as additive and interactive covariate. The x-axis shows chromosome numbers and chromosomal locations (in Mb). **Top panel**, Absolute difference in LOD scores (left y-axis) for each marker. The red horizontal dashed line represents the threshold for significant sex-specific linkage that corresponds to 2 ΔLOD. The boxed inset shows LSM and SE for sucrose intakes of the F_2_ mice grouped by sex and genotype of the markers nearest the linkage peak. Other details are the same as for **Figure 2**.

#### Epistasis QTL

Only for 300 mM sucrose intake (but not for other sucrose consumption measures), the two-dimensional QTL scan identified a significant interaction between markers of on Chr1 (*rs3714728*, 9.0 Mb; *Scon4*) and Chr9 (*rs4227916*, 108 Mb; *Scon3*; **Figure 5A**). This epistatic interaction was confirmed by a generalized linear model analysis (**Figure 5B**).

**Figure 5.**
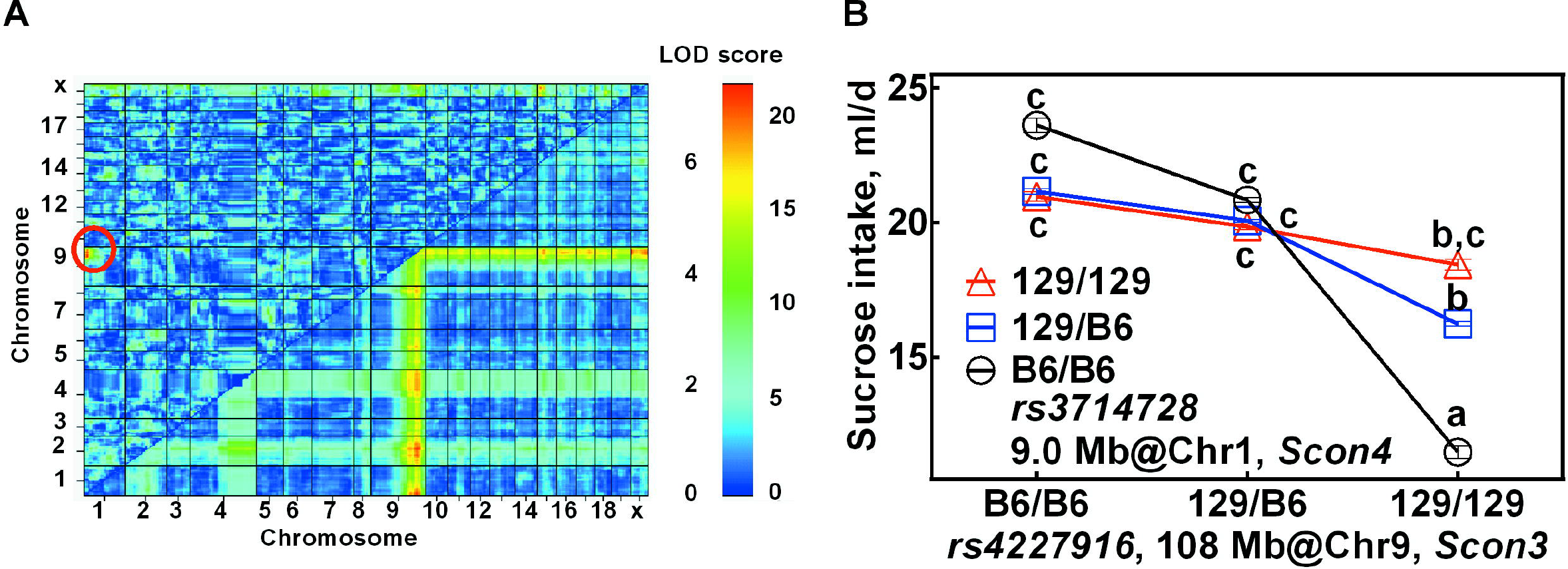
Genome-wide scan of interacting QTLs *Scon3* and *Scon4* for 300 mM sucrose intake. (A) Heat map for a two-dimensional genome scan with a two-QTL model. The maximum LOD score for the full model (two QTLs plus an interaction) is indicated in the lower right triangle. The maximum LOD score for the interaction model is indicated in the upper left triangle. A color-coded scale displays values for the interaction model (LOD threshold = 6.2) and the full model (LOD threshold = 9.1) on the left and right, respectively. A red circle in the upper left section shows significant interaction (LOD = 7.01) between QTLs *Scon3* on Chr9 (108 Mb) and *Scon4* on Chr1 (9 Mb). (B) LSM ± SE for sucrose intakes of the F_2_ mice grouped by genotypes of the markers with the highest epistatic interaction (*rs3714728* and *rs4227916*). The letters (a, b, c), when different, indicate statistically significant differences between genotypes (Tukey’s HSD test: p<0.05; for details, see the Methods section).

#### QTL for the electrophysiological taste responses to sucrose

A QTL for the peripheral neural responses to 100-1000 mM sucrose solutions was mapped to 155.2-156.5 Mb on Chr4 (*Scon2*, corresponding to the *Sac* locus and the *Tas1r3* gene), with the B6 allele increasing the phenotype (**Figure 6**). This locus explained 31-69% of the genetic variance, depending on sucrose concentration (**S7 Table**). There was no effect for lower concentrations (3 and 30 mM) of sucrose (**S7 Figure**).

**Figure 6.**
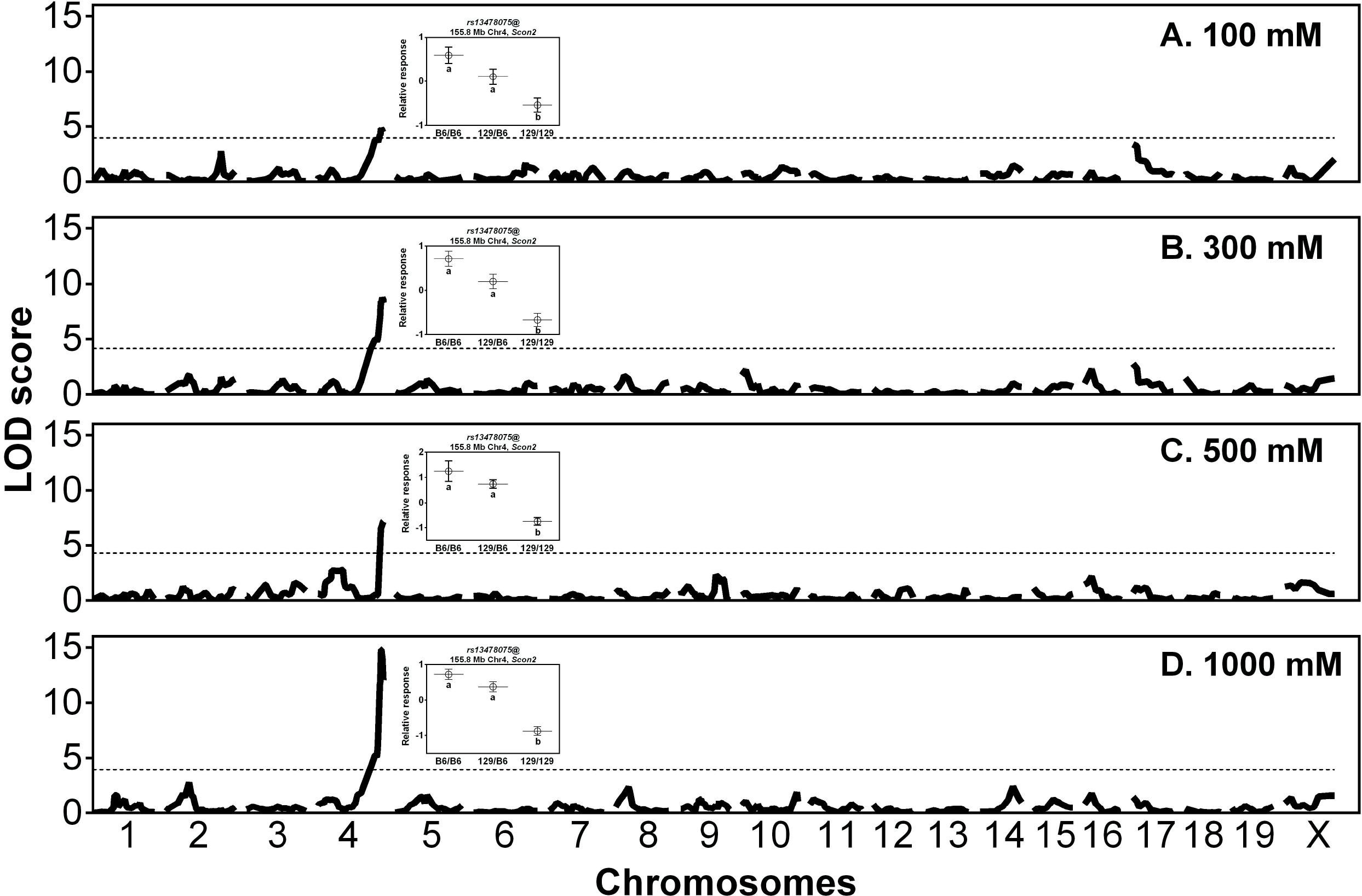
Genome-wide scan of main-effect QTL for peripheral taste responses to sucrose. The horizontal dashed lines represent significant linkage thresholds that correspond to an LOD of 3.96, 4.17, 4.27, and 3.95 for sucrose concentrations of 100 mM (**A**), 300 mM (**B**), 500 mM (**C**), and 1000 mM (**D**), respectively. Boxed insets show means and standard errors for normalized integrated chorda tympani nerve responses to oral stimulation with sucrose of the F_2_ mice grouped by genotype of the markers nearest the linkage peak. Other details are as for Figure 2.

### Candidate genes and their targeted variants

Within the seven critical chromosome regions (*Scon4* on chr1:1-19023602, *Scon6* on chr2:100122595-128989913, *Scon2* on chr4:154415509-1558633334, *Scon7* on chr5:22747915-66662082, *Scon8* on chr8:72545028-129008800, *Scon3* on chr9:87885544-110762590, and *Scon9* on chr15: 40369844-71152967), there are 2565 genes (**S8 Table**), including the taste receptor gene *Tas1r3* in the Chr4 QTL region, known to be associated with behavioral and neural sweet taste responses to sucrose (Inoue, Reed et al. 2004, Bachmanov, Bosak et al. 2011). In total, 1373 missense and stop codon gain or loss variants between parental strains B6 and 129 (**S8 Table**) are within these genes, which include six missense variants, *rs32766606, rs13478071, rs13459081, rs13478079, rs13478082*, and *rs13478081*, within the sweet taste receptor gene *Tas1r3*. All these genes are grouped based on their molecular functions, biological processes, cellular components, protein classes and pathways, and more than 30% genes are categorized into catalytic activity (GO:0003824), about 30% categorized into metabolic process (GO: 0008152), about 17% categorized into cellular membrane (GO: 0016020), about 2% categorized into membrane traffic protein (PC00150), and about 3% categorized into opioid related pathways (P05915, P05916, P05917), and 3% categorized into oxytocin receptor mediated signaling pathway (P04391) (**S9 Table**).

Using results from the two-dimensional scan, we determined if there were candidate genes from the interacting regions that had the same or similar biological function, reasoning that if the two genes have epistatic interaction, they likely act in the same pathway to regulate sucrose intake. We identified 15 gene pairs with the association confidence scores > 0.9 and that contained one gene from the *Scon4 (*on Chr1) QTL region and another from the *Scon3* (on Chr9) QTL region (**S8 Figure, S10 Table**). The interaction of these gene pairs supported the presence of an epistatic interaction effect on mouse sucrose intake.

Gene expression profiling of three mouse tissue sites (striatum, brown and white fat) revealed differential expression of candidate genes (stratum: *n*=15; brown fat: *n*=37, white fat: *n*=171) within the QTL regions (**S11 Table**) between 129 and B6 inbred mouse strains (twofold change with FDR<0.05); the top differentially expressed genes are shown in **Figure 7**.

**Figure 7.**
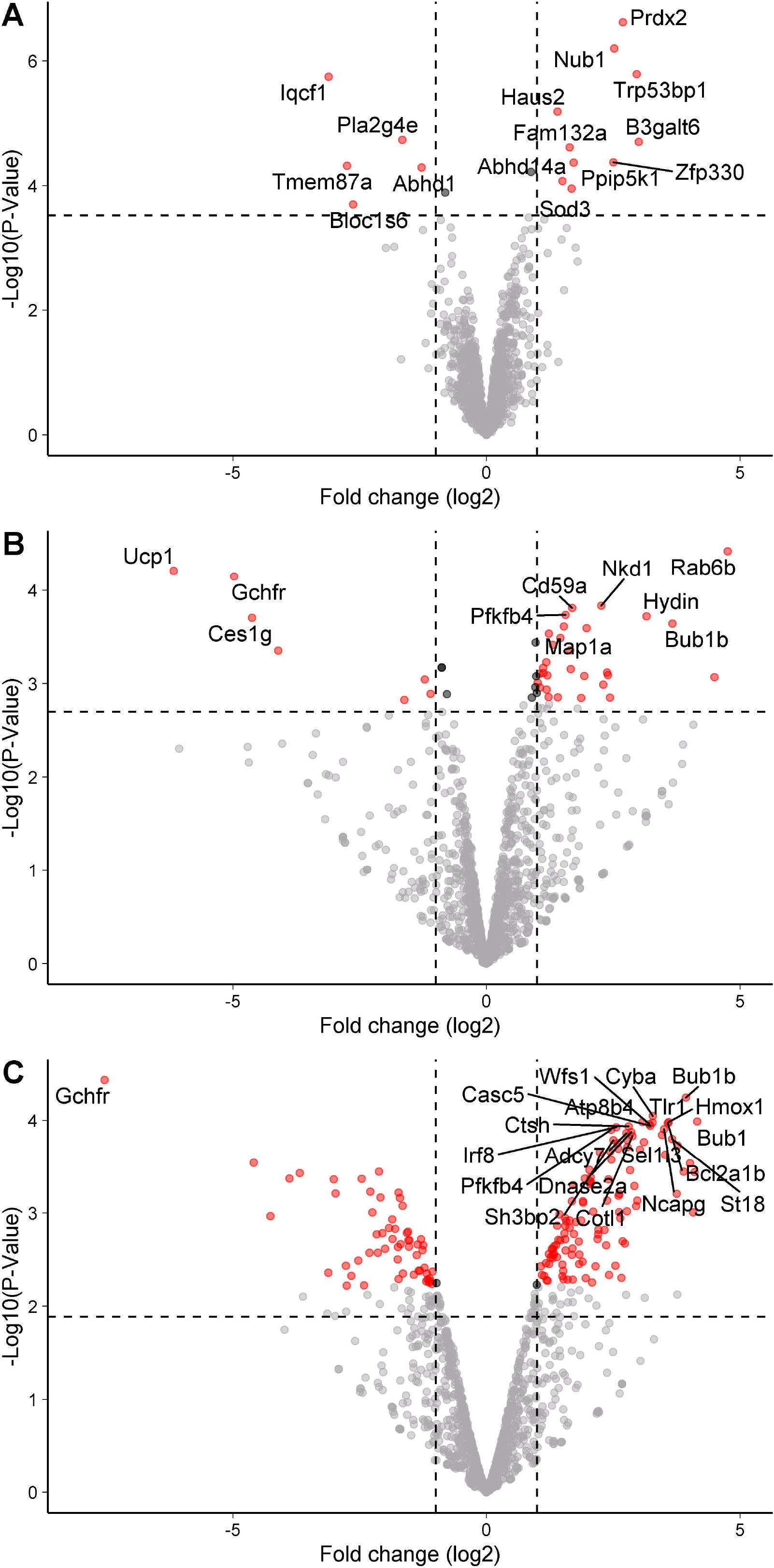
Gene expression profiling shows differential expression of candidate genes (*n*=2675) within the QTL regions between 129 and B6 inbred mouse strains. Microarray-based gene expression analysis data are shown for mouse striatum (**A**), and bulk RNASeq analyses for brown (**B**) and white (**C**) fat. Red dots show differentially expressed genes (twofold changes with FDR<0.05) between two inbred strains, but only the top 10 genes are labeled in B and C. The horizontal dash lines show the corrected significance threshold (FDR=0.05).

## Discussion

In this study, we identified a complex genetic architecture underlying sucrose intake consisting of seven loci, including one sex-specific locus and an epistatically interacting locus pair (*Scon3* and *Scon4*). One locus (*Scon2*) on distal Chr4 harbors the *Tas1r3* gene, which codes for sweet taste receptor subunit T1R3, a major component contributing to behavioral and neural responses to sucrose in mice (Inoue, Reed et al. 2004, Reed, Li et al. 2004). The other loci had significant effects on sucrose consumption, but they apparently do not act at the peripheral level because they did not affect chorda tympani electrophysiological taste responses to sucrose. Several hormones and neuromodulators act as peripheral sweet taste modulators, such as leptin (Kawai, Sugimoto et al. 2000, Nakamura, Sanematsu et al. 2008), oxytocin (Sinclair, Perea-Martinez et al. 2010), glucagon-like peptide-1 (Shin, Martin et al. 2008), cholecystokinin (Hajnal, Covasa et al. 2005), and the central nervous system peptides orexin neuropeptide Y, and opioids (Yirmiya, Lieblich et al. 1988, Marks-Kaufman, Hamm et al. 1989, Dym, Pinhas et al. 2007, Olszewski and Levine 2007), as well as the dopamine D2 receptor (Bulwa, Sharlin et al., Schneider 1989, Dym, Pinhas et al. 2009, Eny, Corey et al. 2009). Furthermore, sucrose intake is controlled by mechanisms beyond the peripheral taste for caloric and sweet substances (Sclafani 2001, de Araujo, Oliveira-Maia et al. 2008, de Araujo 2012); for instance, fibroblast growth factor 21 mediates endocrine control of sugar intake through a hormonal liver-to-brain feedback loop (von Holstein-Rathlou, BonDurant et al. 2016, Soberg, Sandholt et al. 2017) independent of peripheral taste (Dushay, Toschi et al. 2015). Except for *Tas1r3*, none of these genes is located in the seven sucrose QTL regions, so they cannot account for their contribution to sucrose consumption.

We revealed the male-specific QTL (*Scon9*) on Chr15 that affects concentrated sucrose intake (**Figure 1**). Several studies in mice and other species point to sex-specific QTLs (Nuzhdin, Pasyukova et al. 1997, Lionikas, Blizard et al. 2003, Jerez-Timaure, Kearney et al. 2004). For instance, there are male-specific QTLs for alcohol consumption (Melo, Shendure et al. 1996, Peirce, Derr et al. 1998), a sex-specific effect on memory performance when rats drink 10% sucrose daily (Abbott, Morris et al. 2016), and a female-specific QTL (*Dpml*; chr15:51141298-76910293) for dopamine loss in the neostriatum (which serves as a marker of nigrostriatal dysfunction) (Sedelis, Hofele et al. 2003).

In the genome scan for peripheral neural taste responses to sucrose using F_2_ mice, we detected QTL only to distal Chr4 (*Scon2;* encompassing the sweet taste receptor gene *Tas1r3*). However, but we cannot rule out linkages for the neural taste responses to sucrose to other chromosomes: compared with number of mice used for sucrose consumption phenotyping, we measured neural responses in a much smaller number of F_2_ mice, and so there may be lack of statistical power to detect QTLs with smaller effect sizes in other genome regions. We conclude that *Tas1r3* is a major factor contributing to the peripheral taste responses to sucrose and that other sucrose consumption loci are likely independent of peripheral sweet taste.

*Slc5a1* (solute carrier family 5, sodium/glucose cotransporter, member 1), harbored in the *Scon7* (Chr5) QTL region, is expressed in the taste cells and may serve as a mediator of the T1R-independent sweet taste of sugars (Yee, Sukumaran et al. 2011). Thus, *Slc5a1* can be considered a candidate gene for *Scon7*. However, there is no significant linkage to this QTL region for peripheral taste responses to sucrose, suggesting that *Slc5a1* is unlikely a peripheral taste contributor to sucrose intake. Nevertheless, it is possible that we did not detect linkage of the neural responses to sucrose to *Scon7* because of the lack of statistical power. We did not observe any missense variants within this gene between the 129P2/OlaHsd and C57BL/6J strains (closely related to the 129 and B6 parental strains used in our study), but it is possible that there are functional variants in *Slc5a1* between the C57BL/6ByJ and 129P3/J strains that are absent between the 129P2/OlaHsd and C57BL/6J strains (that are not genetically identical with the 129P3/J and C57BL/6ByJ strains).

Excess sugar consumption is a public health concern, which is indicated as a major factor for such human diseases as cardiovascular disease, fatty liver disease, metabolic syndrome, obesity, and diabetes (Ruff, Suchy et al. 2013). This study identified several novel loci that control sucrose intake, which may point to novel pathways beyond peripheral taste sensitivity that could help in understanding the genes involved in sucrose’s effects in other model systems (May, Vaziri et al. 2019), and we may be able to harness these novel mechanisms to control obesity and diabetes and to complement similar efforts to understand the genetic effects on human sucrose intake and preference (von Holstein-Rathlou, BonDurant et al. 2016, Soberg, Sandholt et al. 2017, Hwang, Lin et al. 2019).

## Acknowledgments

We gratefully acknowledge Maria L. Theodorides, Zakiyyah Smith, Mauricio Avigdor, and Amy Colihan for assistance with animal breeding. We also acknowledge Richard Copeland and the consistent high-quality assistance of the animal care staff at the Monell Chemical Senses Center and thank them for their service. Yutaka Ishiwatari assisted with genotyping markers.

## DECLARATIONS

### Ethics approval

All animal procedures in this study were approved by the Institutional Care and Use Committee of the Monell Chemical Senses Center.

### Consent to participate

Not applicable.

### Consent for publication

Not applicable.

**Availability of data and material:**

### Code availability

Not applicable.

**Accession IDs:**

### Author contributions

AB and DR designed the study. CL conducted the study. CL and DR analyzed data. MI conduct electrophysiological experiment. XL and NB genotyped mice. CL and DR wrote the paper. AB, MT and GB commented and edited the paper. All authors read the paper and approved its contents.

## Figure legends

**S1 Figure. Imputation of missing data of behavioral phenotypes**

(**A**) Missing data portions of each trait for intake (Int) and preference (Pre) of 3, 120, or 300 mM sucrose. (**B**) Summary of missing phenotype data pattern: red, missing data; blue, observed data. (**C**) Plausible values for the imputed data points (magenta; imputed by multiple imputation by chained equations) and observed data points (blue). The x-axis shows imputation numbers: 0 = observed data; 1-5 = data of 1-5 imputations. The y-axis shows intake (ml) or preference (%).

**S2 Figure. Distribution of sucrose intake and preference in female and male F**_**2**_ **mice**

The y-axis shows density, the fraction of number of animals with sucrose intake or preference (x-axis). Colors show different sucrose concentrations, and the white dashed lines indicate means for each concentration. **Top row**, Unadjusted sucrose intake and preference data. **Bottom row**, Sucrose intake residual scores standardized to habitual water intakes calculated within each sex, and sucrose preferences standardized within sex and experiment.

**S3 Figure. Dominance effects for 3 mM sucrose intake and preference by sex**

Data are mean ± SE for sucrose solution intake (left) and preference (right) by male and female parental strains (B6, 129) and F_2_ mice. Values for the F_2_ mice (data points with center bar) are displayed at the parent midpoint to show the direction of dominance, if any. Asterisks indicate a dominant mode of inheritance (as defined by a significant difference between the F_2_ group average and the parental midpoint).

**S4 Figure. Heat map of Pearson correlations for sucrose intakes and preference scores of the F**_**2**_ **mice (*N*=623)**

Missing and imputed data for each trait are summarized in **S1 Figure**. For correlation analyses shown here, we used the first imputation (imputation number 1 in S1 Figure).

**S5 Figure. Correlations between sucrose intake (ml/d) and phenotypes of F**_**2**_ **mice**

Sucrose intake data are shown for unadjusted data (blue) and adjusted residual scores (red). Phenotypes were sex (M, F), total body weight (BW; g), total body fat (g), body length (cm), daily habitual water intake (Wat; ml/d), and age (months). N = 452 for 3 and 300 mM sucrose; N = 623 for 120 mM sucrose.

**S6 Figure. Correlations between sucrose preference (%) and phenotypes of the F**_**2**_ **mice**.

Sucrose preference data are shown for unadjusted (blue) and adjusted standardization scores (red). For details, see **S5 Figure**.

**S7 Figure. Chorda tympani responses to sucrose (relative to 100 mM NH**_**4**_**Cl) in F_2_ mice**

Data are standardized within sex and experiments, grouped by *Tas1r3* genotype. Values are means ± standard errors. *p<0.01, one-way ANOVA.

**S8 Figure. Protein-protein associations for genes within the sucrose epistasis QTL *Scon3* and *Scon4* confidence interval regions**

Protein-protein associations for genes within QTL confidence intervals of *Scon4* (on Chr1) *Scon3* (on Chr9) (see **S11 Table**) identified by searching protein names using information from high-throughput experimental data, mining of literature databases, and predictions based on genomic context analysis.

