## Supplemental Table 1 for "Genetics of mouse behavioral and peripheral neural responses to sucrose"

**S1 Table.** Number of mice used for behavioral and gustatory neural response experiments

| **Experiment** | **Mouse type** | **Behavioral test (M/F)** | **Neural responses test (M/F)** |
| --- | --- | --- | --- |
| Experiment 1^a^ | F_2_ | 171 | 19 |
| Experiment 2 | F_2_ | 228/228 | 29/29 |
| Experiment 2 | 129P3/J | 10/10 | Not available |
| Experiment 2 | C57BL/6ByJ | 10/10 | Not available |
| Additional test | F_2_ | Not available | 2/5 |

M=males; F=females.

^a^Only male mice were used in Experiment 1.
